# Delays, behaviour and transmission: modelling the impact of effective private provider engagement on tuberculosis control in urban India

**DOI:** 10.1101/461426

**Authors:** Nimalan Arinaminpathy, Sarang Deo, Simrita Singh, Sunil Khaparde, Raghuram Rao, Bhavin Vadera, Niraj Kulshrestha, Devesh Gupta, Kiran Rade, Sreenivas Achuthan Nair, Puneet Dewan

## Abstract

In India, the country with the world’s largest burden of tuberculosis (TB), most patients first seek care in the private healthcare sector, which is fragmented and unregulated. Ongoing initiatives are demonstrating effective approaches for engaging with this sector, and form a central part of India’s recent National Strategic Plan: here we aimed to address their potential impact on TB transmission in urban settings, when taken to scale. We developed a mathematical model of TB transmission dynamics, calibrated to urban populations in Mumbai and Patna, two major cities in India where pilot interventions are currently ongoing.

We found that, when taken to sufficient scale to capture 75% of patient-provider interactions, the intervention could reduce incidence by upto 21.3% (95% Bayesian credible interval (CrI) 13.0 – 32.5%) and 15.8% (95% CrI 7.8 – 28.2%) in Mumbai and Patna respectively, between 2018 and 2025. There is a stronger impact on TB mortality, with a reduction of up to 38.1% (95% CrI 20.0 – 55.1%) in the example of Mumbai. The incidence impact of this intervention alone may be limited by the amount of transmission that has already occurred by the time a patient first presents for care: model estimates suggest an initial patient delay of 4-5 months before first seeking care, followed by a diagnostic delay of 1-2 months before ultimately initiating TB treatment. Our results suggest that the transmission impact of such interventions could be maximised by additional measures to encourage early uptake of TB services.

---

India has the world’s largest burden of tuberculosis (TB) ^1^. Over the past two decades India’s Revised National Tuberculosis Control Programme (RNTCP) has made notable progress in reducing TB deaths, through the provision of basic TB services via the public sector ^2–5^. Nonetheless, major challenges remain: healthcare in India is dominated by the private sector, where the majority of patients first seek care^6–9^. Private healthcare providers often use inaccurate diagnostic tests for TB, or omit testing altogether, leading to diagnostic delays while patients cycle between different providers ^7,10,11^. Even once patients are diagnosed, a general lack of treatment adherence monitoring and support is unfavourable for long-term treatment outcomes^12^. Moreover, although tuberculosis was made a notifiable disease in 2012 ^13^, there remain major challenges in encouraging private providers to comply with these obligations ^14,15^. For these reasons, in India’s recently-announced plan to eliminate TB, private sector engagement forms a key strategic priority ^16^.

In a demonstration of private sector engagement in India, the ‘Public Private Support Agency’ (PPSA) model used a combination of patient subsidies and provider incentives to encourage higher standards of diagnosis and treatment amongst private providers ^17^. Originally implemented in two Indian cities, Mumbai and Patna (respectively by the NGOs PATH and World Health Partners), these measures have yielded rapid increase in TB notification from the private sector ^3^. However, their potential epidemiological impact remains unclear; measuring such impact empirically presents prohibitive challenges in the intervention coverage, population size and study duration that would be needed.

Here we take an alternative approach, using a dynamical model of TB transmission, developed to capture the complexity of careseeking in urban settings in India. The model is calibrated to detailed patient careseeking surveys in Mumbai and Patna, as well as data on TB epidemiology in these settings. While Patna is typical of an urban setting in India, Mumbai is exceptional in its high burden of MDR-TB ^18,19^. We ask: What impact could such engagement have on TB transmission, in particular on TB incidence? What are the key drivers of this impact?

In what follows we present an overview of the model framework, with further details in the supporting information. We describe the pathway surveys, and the approach for incorporating this evidence in the model framework. We then present results for the potential epidemiological impact of private sector engagement in Mumbai and in Patna, followed by an examination of the drivers of this impact: in particular, we investigate specific types of patient and provider behaviour that matter most for TB transmission. Finally we discuss implications for controlling TB transmission in India, and important questions arising for future work.

## Methods

### Model overview

We developed a deterministic, compartmental model, whose overall structure is illustrated schematically in Figure 1. The model divides the population into different states, reflecting their disease and careseeking states, with a set of coupled, differential equations capturing transmission dynamics, and the transitions between states (see appendix). We first give an overview of the essential dynamical processes captured by the model, before describing the evidence sources used to quantify these dynamics.

**Figure 1.**
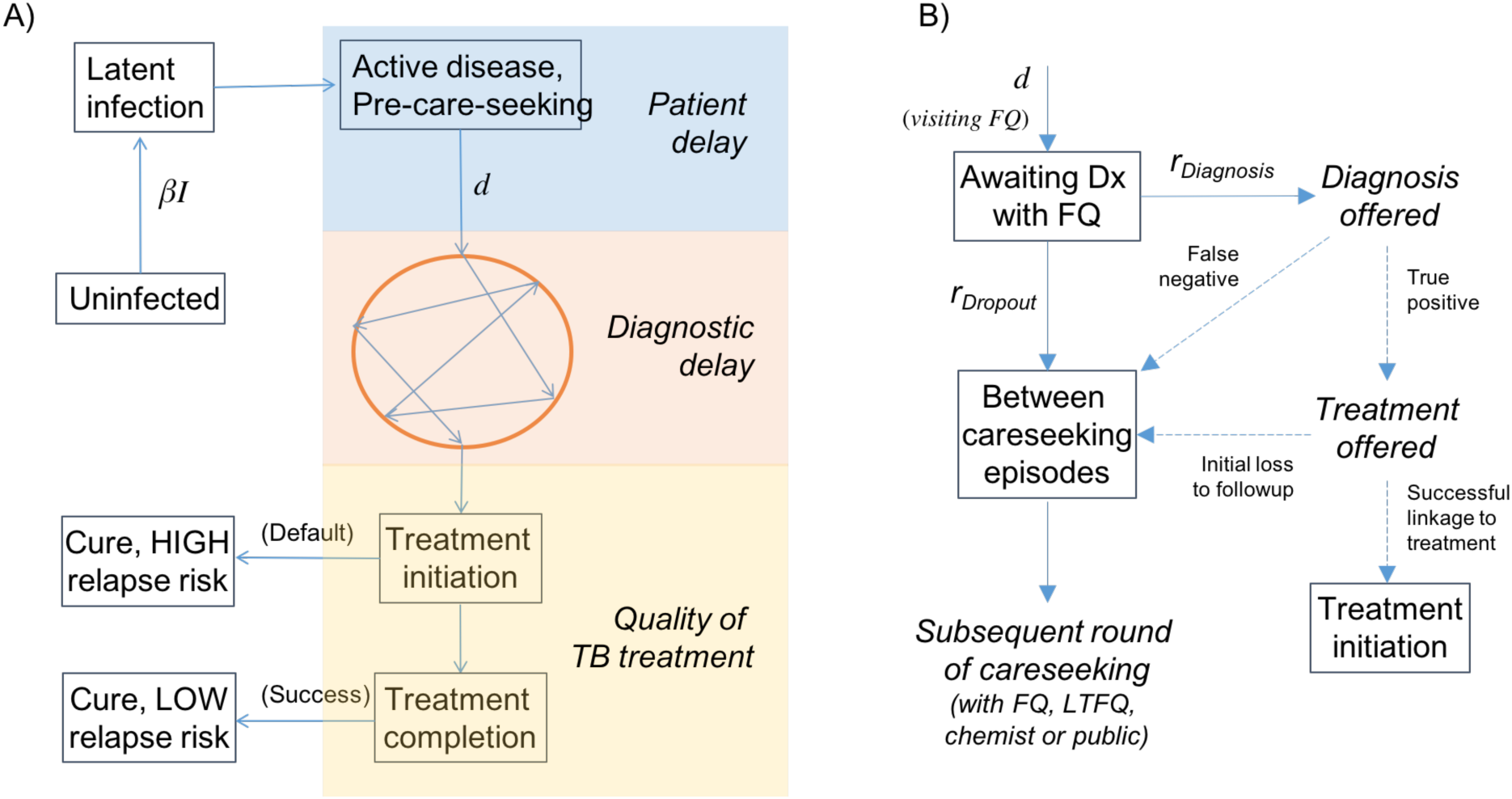
Schematic illustration of the transmission model. (A) The figure shows two important parameters in the model, the annual infections per active TB case (β) and the mean, per-capita rate of careseeking once a patient develops active TB (d), which are calibrated to yield the correct ARTI and prevalence (see Table S2). The ‘bubble’ in orange denotes the sequence of providers that a patient visits before receiving a TB diagnosis. Here, we distinguish the associated ‘diagnostic delay’ with the initial ‘patient delay’. This model also includes the acquisition and transmission of multi-drug-resistant (MDR) TB, not shown here for clarity. (B) Detail of the diagnostic process depicted in the ‘bubble’ in panel (A), showing the case of a formally qualified (FQ) provider (this structure applies also to other provider types). Here and elsewhere, ‘Dx’ denotes ‘diagnosis’. Solid lines represent hazard rates in the model, while dashed lines represent proportions. Note the ‘competing hazards’ of diagnosis vs patient dropout. Terms in boxes represent compartments in the model, while terms in italics show intermediate stages, associated with the quality of TB care (accuracy of diagnosis, and treatment initiation).

We assumed that each active case of TB causes, on average, *β* infections per year. We further assumed that, upon development of active disease, there is a ‘patient delay’ before first seeking care. In the model equations (see supporting information), this delay is governed by the per-capita careseeking rate *d*. As described below, *β* and *d* are calibrated for consistency with the TB epidemiology in urban slums. Once patients enter the careseeking pathway (denoted by the circle in Fig. 1A), they visit a series of providers: the resulting ‘diagnostic delay’ is the interval from first careseeking to initiation of anti-TB treatment. This delay is governed by the timeliness with which these providers can offer an accurate TB diagnosis, and retain a patient for long enough to initiate appropriate treatment.

Upon initiating treatment, patients exit the diagnostic pathway illustrated in Figure 1A, where the next hurdle is to complete high-quality (DOTS standard) treatment. Most patients in the private sector lack adherence support, and thus do not complete the 6-month, first-line regimen ^12,20^: we assume that those defaulting from treatment, although immediately lacking infectiousness and being relieved of symptoms, face an increased risk of relapse in the long term, compared to patients successfully completing the 6-month regimen, with a parameter conservatively sourced from clinical trials of shorter durations of rifampicin treatment.

In this framework, a PPSA has two functions: (i) to subsidise high-quality diagnosis for patients in the private sector, increasing the probability of an accurate TB diagnosis, and thus reducing the overall diagnostic delay (depending on coverage, or the proportion of providers engaged), and (ii) providing adherence support to maximize treatment completion. In both cases, we assumed that private providers engaged by a PPSA are able to match the quality of TB care in the public sector, on these dimensions.

For simplicity we ignored HIV/TB coinfection, which is estimated to account for only 5% and 1% of notified TB cases in Maharashtra and Bihar, respectively ^3^. However, we incorporated the acquisition and transmission of multi-drug-resistant (MDR) TB. In particular, we assumed that each infectious case of MDR-TB, not undergoing appropriate second-line treatment, causes **β*_MDR_* infections per year, to be calibrated to the estimated burden of drug resistance (see below). We assumed that there is essentially no management of MDR-TB in the private sector, and populated parameters for second-line treatment outcomes in the public sector to match those reported by RNTCP ^3^.

### Epidemiological inputs

WHO estimates for incidence and prevalence, although often used to inform transmission models ^21–23^, pose two important limitations for the present work. First, national incidence estimates for India are informed by expert opinion on the proportion of cases that are notified to RNTCP ^24^, which itself is subject to change ^1^. Second, WHO national estimates do not address subnational heterogeneity, and thus would not accurately reflect the epidemiological conditions in urban settings considered in our study.

Instead, to relate the model as closely as possible to the primary data available, we used the Annual Risk of TB Infection (ARTI, a measure of the intensity of transmission in a given setting), and the prevalence of TB, as estimated by subnational prevalence surveys in India. Unfortunately, neither Mumbai nor Patna has yet had a prevalence or infection survey (to inform prevalence or ARTI estimates, respectively). Nonetheless, infection surveys in Chennai and Delhi ^25^ suggest that ARTI in urban settings is in the range of 2–3%. We adopted this range in modelling Mumbai and Patna populations as well. For prevalence, we borrowed from a recent prevalence survey in Chennai, which estimated urban prevalence at 388 cases per 100,000 population ^26^. To accommodate the uncertainty in applying these estimates to settings outside Chennai, As both prevalence and ARTI estimates are being borrowed from other settings, we incorporated broad uncertainty in applying these estimates in the present study. For example, for prevalence estimates we adopted uncertainty intervals 25% wider than those published for Chennai (see table S2, supporting information).

For the burden of drug resistance, we assumed that Patna is typical of the national average, with 3 - 5% of incident TB cases being MDR-TB. For Mumbai, we used program-reported data on routine surveillance for drug-resistant TB to populate a more extreme scenario for drug resistance, assuming that 8 – 16% of incident cases have MDR-TB. These inputs are summarized in table S2, supporting information.

### Patient pathways

We adopted four different categories of provider: (i) those in the public sector (DOTS facilities); (ii) private chemists; (iii) private, ‘fully qualified’ (FQ) providers with qualifications in allopathic medicine; (iv) and private, ‘less-than-fully-qualified’ (LTFQ) providers with other medical qualifications, or none at all.

We used data from community-based patient pathway surveys, recently conducted in Mumbai (76 TB patients and 196 patient-provider interactions) and Patna (64 TB patients and 121 patient-provider interactions), and described in detail elsewhere ^11,27^. In brief, individuals in the community, who had been on TB treatment within the preceding 6 months, were administered an in-depth interview, to identify the sequence and types of providers that each patient visited before their TB diagnosis. Although subject to the usual limitations of patient recall ^28^, this community-based survey has nonetheless cast unprecedented light on the careseeking patterns in these urban slum settings ^11^.

A patient’s contact with a given provider may last several days, sometimes weeks: this process ends either when the provider eventually suspects and confirms TB, or when the patient drops out to visit an alternative provider. Here, we model this combination of behaviours using independent, competing exponential hazards, taking both to be specific to the type of provider involved (public, FQ, LTFQ or chemist). Figure 1B shows the overall framework: for Mumbai and Patna separately, we used the pathway survey data to estimate the hazard rates *r_Diagnosis_, r_Dropout_* in Fig. 1B, as well as the probabilities of accurate diagnosis per provider visit. We also used this data to estimate the role of different provider types in the careseeking pathway, in particular: the proportions of patients visiting each type of provider on the first careseeking attempt, and the corresponding proportions on subsequent visits, conditional on the type of provider last seen. We used the Expectation-Maximisation algorithm as a systematic approach for estimating rates and uncertainty (see supporting information for further details).

For parameters related to the treatment cascade (the proportion of TB diagnoses initiating and completing treatment), we draw from a recent systematic review for the public sector ^29^. In the absence of systematic evidence for private providers, we incorporate plausible uncertainty distributions for these parameters (Table S2, supporting information).

### Simulating impact

In both Mumbai and Patna, evidence suggests a marked heterogeneity amongst providers, with certain specialist providers handling a substantially higher TB caseload than others. While this suggests important opportunities for efficiency, by ‘targeting’ such providers, in the present work, for simplicity we chose instead to measure PPSA ‘coverage’ from a patient perspective: that is, the proportion of patient-provider interactions that involve a PPSA-engaged provider. Thus, for example, we present a 75% ‘coverage’ in the understanding that – in practice – this could be brought about by recruiting fewer than 75% of providers, in a targeted way.

For a given PPSA coverage, we simulated cumulative TB incidence and mortality between 2018 and 2025. We then estimated the TB cases and deaths averted, relative to a ‘no-PPSA’ baseline, with the standard of TB care in public and private sectors projected forward without change. We simulated two types of PPSA: an ‘accurate diagnostic’ scenario in which engaged providers have diagnostic accuracy equal to those of the public sector, and a ‘timely diagnostic’ scenario which, as well as accurate diagnosis, additionally encouraged private providers to conduct a diagnostic test as early as possible (whether for TB or not). Note that, in both cases, treatment outcomes were also assumed to be improved to the level of the public sector.

### Uncertainty

We used a Bayesian melding procedure ^30^ to capture uncertainty in the epidemiological and pathway inputs described above, as well as in other input parameters in the model (see uncertainty ranges in tables S2, S3, supporting information). In brief, this procedure yields 100,000 parameter sets that, in ensemble, capture simultaneously the uncertainty in the parameter inputs, and in the data. Projecting the epidemiological impact of a PPSA from each of these parameter sets, under given scenarios for PPSA coverage, we then calculated the central estimate and uncertainty in impact by calculating the 2.5^th^, 50^th^ and 97.5^th^ percentiles in the outcomes of interest (lives saved, percent cases averted). We refer to these uncertainty intervals as ‘credible intervals’ (CrI) to distinguish them from the ‘confidence intervals’ arising from frequentist statistical approaches. Further details are provided in the supporting information.

The model includes several different parameters (including epidemiological inputs). To identify those parameters that are most important for model findings, we performed a multivariate sensitivity analysis on the output of the Bayesian analysis described above. In particular, we examined which model inputs accounted for the greatest amount of uncertainty in model outputs: that is, the inputs that are most influential in the precision of the model output. To do this we selected, as a model output, the percent cases averted by a PPSA intervention at 75% coverage under the ‘timely diagnostic’ scenario described above, in both cities. We computed the partial rank correlation coefficient (PRCC) between this output and each of the model parameters: in brief, the PRCC quantifies the correlation between a given model input and the model output, when variance in all other parameters has been accounted for. Those model inputs expressing the greatest PRCC are those to which the model is most sensitive.

As well as this parameter uncertainty, we additionally tested the model sensitivity to two forms of structural uncertainty: (i) First, in the simulations described above we assumed that each TB case undergoes a constant infectiousness *β* through time. In practice, over time **β** may increase (for example if bacillary load rises with symptom severity), or decrease (for example if TB patients exhaust their closest contacts as opportunities for infection), with implications for the transmission that a PPSA could impact ^31^. To capture these scenarios in a simple way, we assumed that infectivity during the patient delay in Figure 1A is *k* times that during the diagnostic delay. We tested sensitivity of model findings to *k*. (ii) Second, the PPSA we have modelled is a combination of interventions, each involving different indicators for the quality of TB care in the private sector. To examine the most important, we simulated a ‘partial’ PPSA that could implement improvements in all but one of the indicators for quality of care. We recorded the resulting drop in impact (percent cases averted), relative to a ‘full’ PPSA, and repeated this analysis for each of the indicators involved.

## Results

Figure 2 shows the model fits for prevalence, ARTI and percent MDR-TB, in both cities. The sampled parameters show agreement with the estimates in ARTI and prevalence data, while also accommodating the range of uncertainty in these inputs. Estimated parameter values are shown in Tables S3, S4, supporting information. Mumbai and Patna show contrasting careseeking patterns, as illustrated by Figure S1 in the appendix (see also Table S3). For example, chemists play a stronger role in first careseeking in Patna than in Mumbai, while for formally qualified providers the converse is the case. These differences underline the potential heterogeneity in healthcare settings across India.

**Figure 2.**
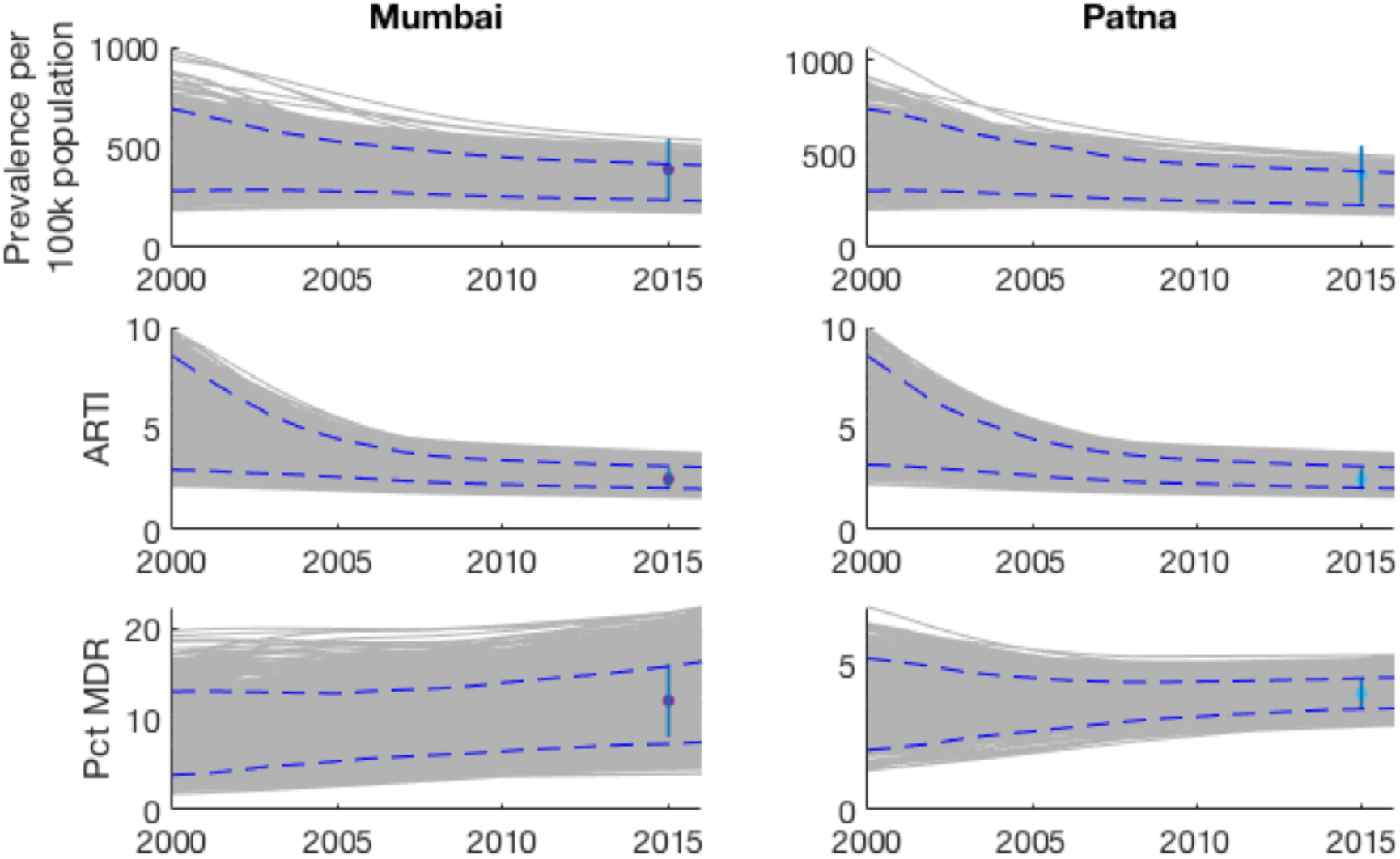
Illustration of the model fits to key epidemiological indicators. (Prevalence, ARTI and proportion of TB cases being multi-drug resistant). Shown are the epidemic trajectories corresponding to each of the sample parameter sets (in grey); the simulated 95% credible intervals over time (blue dashed lines); and the calibration targets (points and vertical ranges, plotted in 2015).

Figure 3 illustrates the potential epidemiological impact of a PPSA in Mumbai, assuming an intervention that scales up over 5 years from 2018 to cover 75% of patient visits to a provider. Such an intervention is focused on improving diagnostic accuracy and treatment outcomes in the private sector, without addressing the promptness with which a provider offers a diagnosis. A PPSA of this scale would reduce cumulative TB incidence by 8.5% (95% CrI 4.2 – 15.6%) over the next ten years. There is a stronger impact on MDR-TB, with a reduction of 21.2% (95% CrI 13.0 – 32.5%) in cumulative incidence. Further, a PPSA of this scale could have a substantial effect on TB mortality, reducing TB deaths by 21.7% (95% CrI 10.6 – 35.0%).

**Figure 3.**
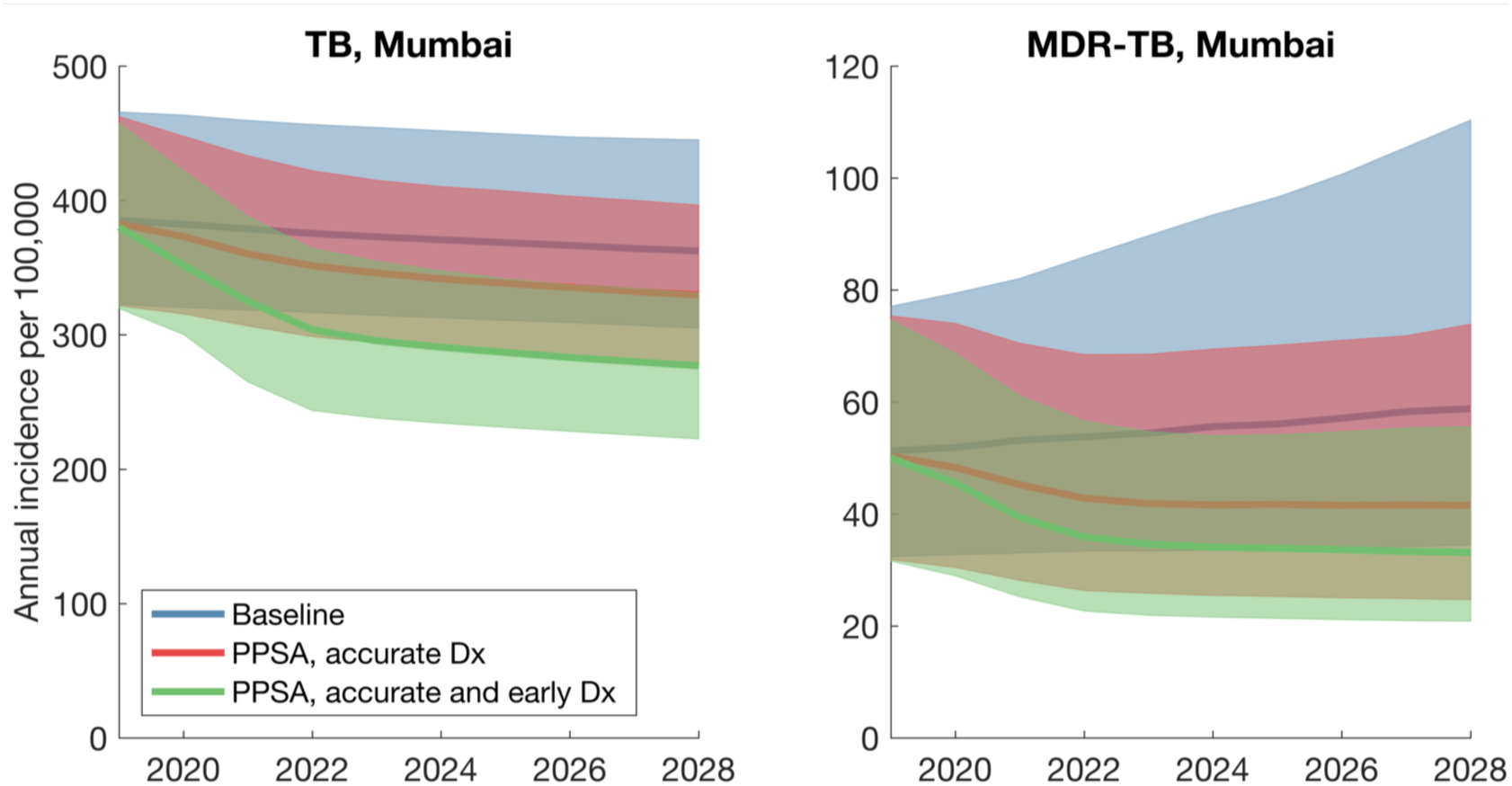
**Illustration of the TB dynamics under scale-up of a PPSA**, in the example of Mumbai. These results capture the scenario of a PPSA being scaled up (over three years from 2018) to cover 75% of patient-provider interactions. Lines show central estimates, and shaded regions show 95% credible intervals.

If providers are additionally encouraged to order a diagnostic test as early as possible (i.e. a ‘timely diagnosis’ scenario to pre-empt patient dropout), PPSA impact increases substantially, to an incidence reduction of 21.4% (95% CrI 11.1 – 32.7%) and a mortality reduction of 38.1% (95% CrI 20.0 – 55.1%). Figure S2 (supporting information) shows similar, corresponding results for Patna. Figure S3 (supporting information) illustrates how these types of impact could vary with PPSA coverage.

To examine factors that may be limiting the impact shown in Fig. 3, we examined the model estimates for the patient and diagnostic delays illustrated in Fig. 1. As illustrated in Fig. 4, while the simulated diagnostic delay is consistent with the 1 month estimated in previous analysis ^8,11^, results suggest that the initial patient delay could be still longer, at 4.4 months and 5.2 months in Mumbai and Patna, respectively, although with broad uncertainty around these estimates. Figure S4 in the appendix shows the potential epidemiological impact of a PPSA that is enhanced by measures to shorten the patient delay; below we discuss possible examples of such measures.

**Figure 4.**
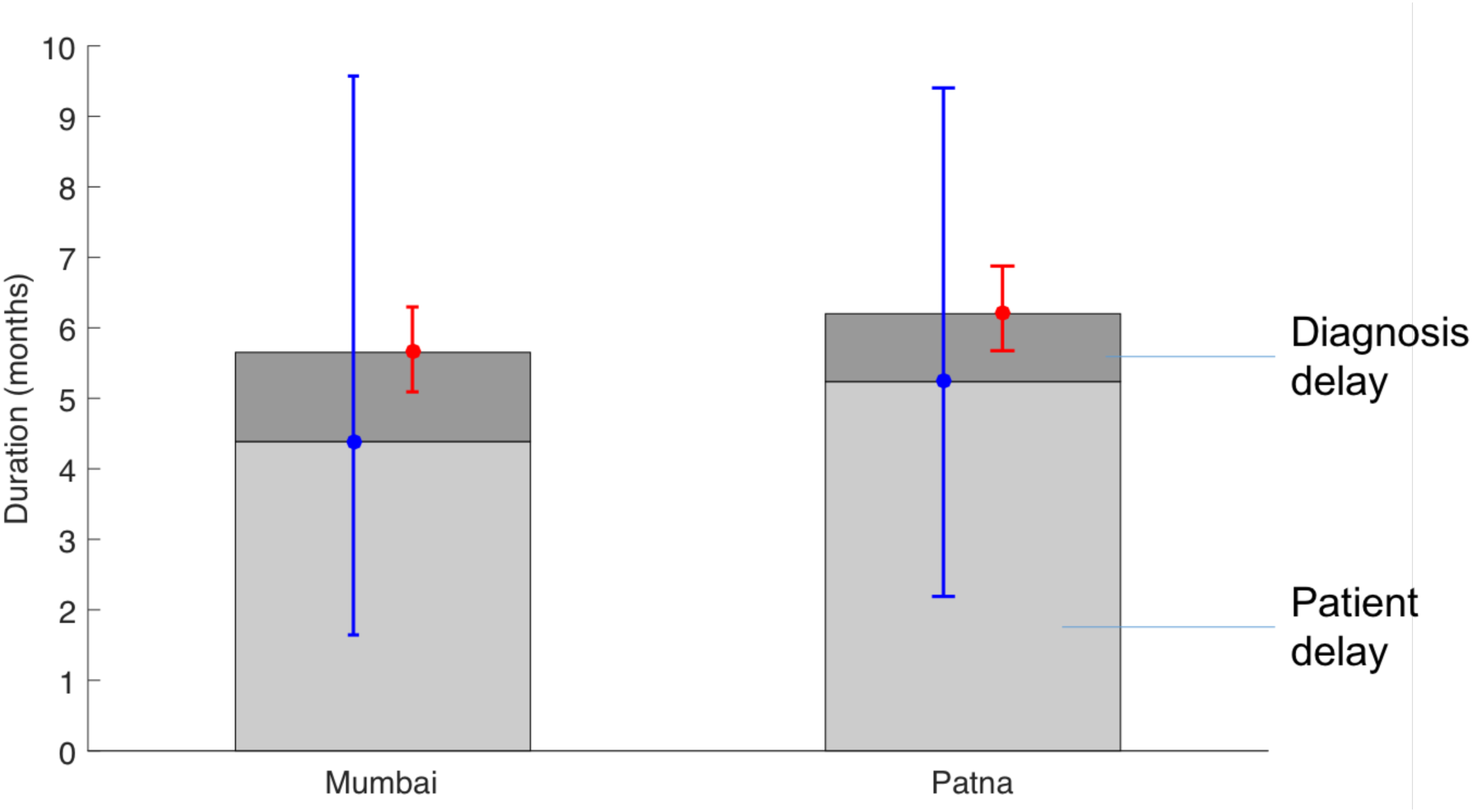
**Components of the mean infectious period**, i.e. the duration from the start of active disease to treatment initiation, death or self cure. Simulated in the absence of any PPSA intervention. The light grey region shows the simulated patient delay, while the dark grey region shows the delay in diagnosis (i.e. from first provider visit). Error bars in blue and red show the uncertainty in these estimates, respectively. The patient delay estimate is driven by prevalence and ARTI, while the diagnostic delay estimate is driven by the process illustrated in Fig.1B. A PPSA addressing only patient behaviour would impact only the dark grey regions.

Figure 5 shows the results of parameter sensitivity analysis, in which we quantified the influence of each model input against ‘simulated impact’, the latter measured as the percent cases averted by a PPSA at 75% coverage in both Mumbai and Patna (corresponding to the green shaded region in Figure 3). Figure 5 illustrates the importance of epidemiological inputs, for this output. In both cities, the assumed prevalence and ARTI are the model inputs accounting for the greatest amount of output uncertainty. Where the true value of prevalence in either city lies towards the lower end of the assumed range, the percent cases averted approaches the upper end of the uncertainty illustrated in Figure 3, and vice versa for ARTI. In both settings the levels initial loss to followup in the public sector (i.e. those diagnosed who do not initiate treatment) is also a leading factor; remaining parameters, to which the model is less sensitive, depend on the local conditions in both of these settings.

**Figure 5.**
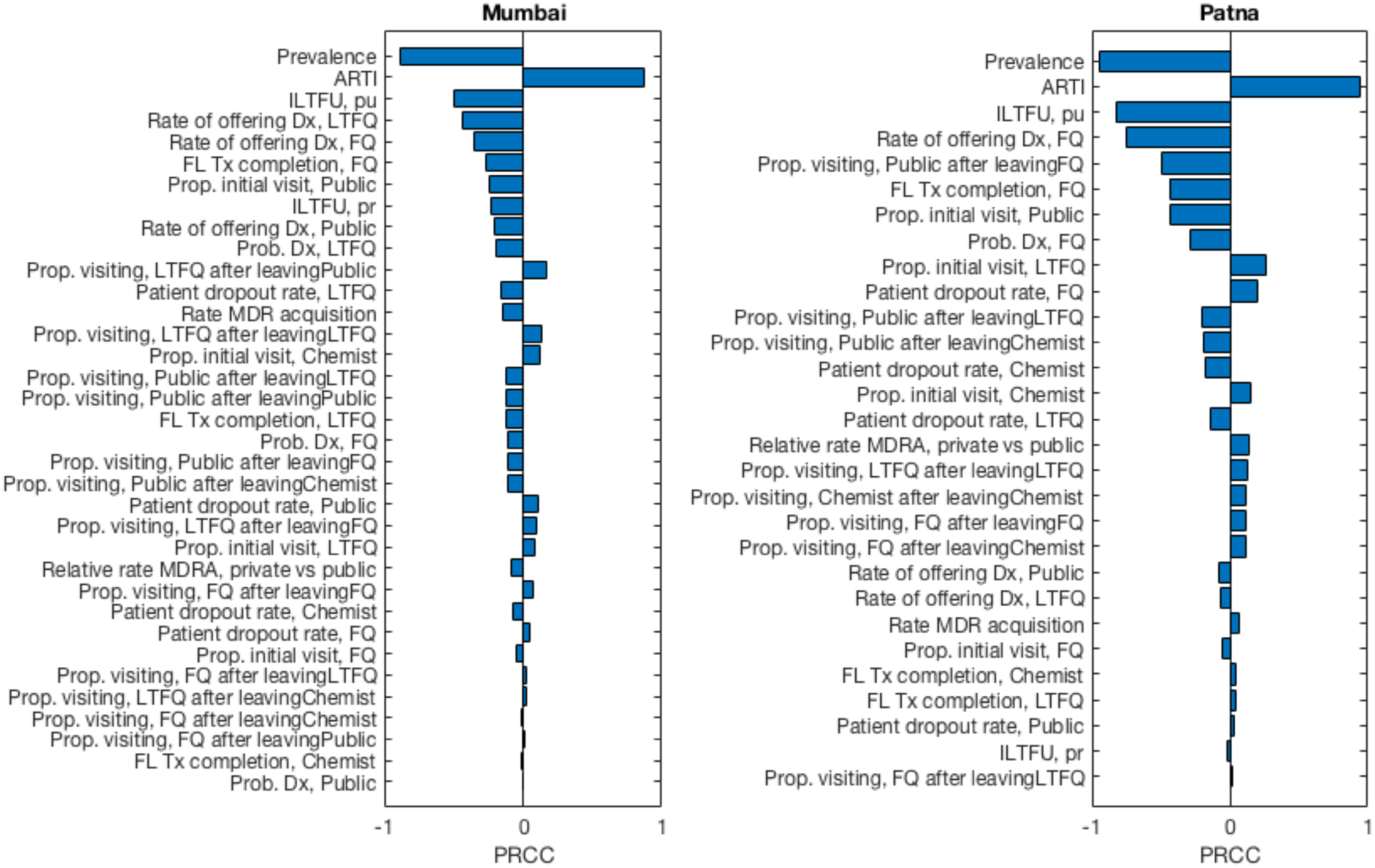
Multivariate sensitivity analysis of model inputs (parameters and data). Bars show the partial rank correlation coefficient (PRCC) between each model input and a selected output: ‘simulated impact’, or the percent cases averted by a PPSA acting at 75% coverage, with accurate and early diagnosis. Figures show that in both Mumbai and Patna, the two model inputs to which simulated impact is most sensitive are the prevalence and ARTI. Prevalence has a negative partial rank correlation with impact: that is, lower values of prevalence are associated with higher levels of impact, and vice versa for ARTI. Note that the combined effect of uncertainty in all of these parameters corresponds to the full uncertainty range illustrated in Figure 3A, green shaded region. This range indicates the maximum extent to which model outputs could diverge from central estimates, subject to the assumed uncertainty ranges in model inputs.

In addition to addressing parameter uncertainty we finally conducted sensitivity analysis to some underlying assumptions. First, as described above, we allowed for differential infectiousness in the two stages of delay shown in Figure 4. Figure 6A shows results for the percent cases averted, as a function of the longitudinal variation in infectiousness. As expected, scenarios with increasing infectivity over a patient’s clinical course (decreasing *k* in the figure) yield greater predicted impact of a PPSA.

**Figure 6.**
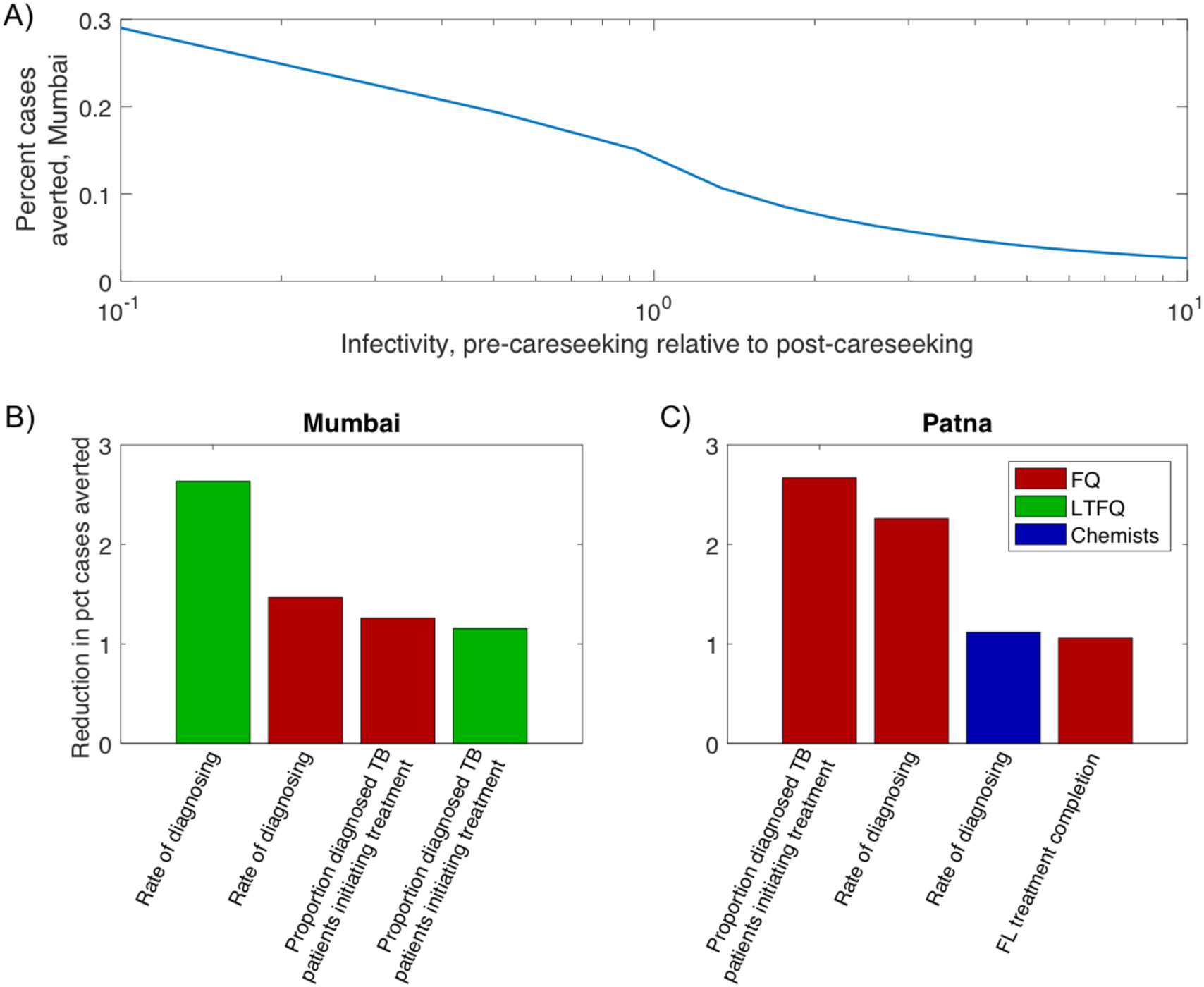
Sensitivity analysis for key assumptions in the model. (A) Effect of assumptions for how TB infectivity varies during the clinical course. Shown is the impact of a PPSA at 75% coverage in Mumbai (percent cases averted over ten years). The x-axis shows a range of scenarios for the infectivity during the patient delay, relative to that during the diagnostic delay. (B, C) Identifying key elements of private provider behaviour. The figures show the drop in overall impact that results, when a PPSA that fails to improve the provider behaviour shown (while addressing all remaining provider behaviours). For clarity, plots show only the four most important factors in each setting. Bar colours denote different provider types, as shown in the right-hand legen

Second, we examined the sensitivity of projected impact to the assumption that all PPSA activities are performed effectively. We aimed to identify which activities accounted for most of the impact shown in Figure 3. Results, shown in Figs. 6 B – C, suggest that in Mumbai, the quality of diagnosis and treatment amongst LTFQ providers is key. In Patna, by contrast, the quality of care amongst FQ providers is most important. Echoing the contrasting pathways illustrated in Figure S1, these results highlight how intervention priorities in different cities may need to be tailored to the local conditions.

## Discussion

Engaging with India’s vast, fragmented private healthcare sector is a key step in enhancing TB control in India. Our work adds to other modelling studies capturing the role of the private sector in TB care in India, including a multi-model comparison examining packages of interventions in the context of the End TB goals ^23^, and the potential impact of implementing molecular diagnostics in the private sector ^21^. A strength of the current work is that it is informed by unique, detailed patient pathway data from Mumbai and Patna. This data enables us to analyse the relative importance of the different delays illustrated in Figure 1, to a greater extent than in previous work.

Our findings illustrate that a PPSA taken to scale in urban settings, such as Mumbai and Patna, could have a meaningful impact on TB burden (Fig.3, Fig.S3). Improved diagnosis and treatment adherence could strongly reduce TB mortality. Moreover, the use of rapid molecular tests in the private sector could have strong implications for MDR-TB: by facilitating the early recognition of drug sensitivity status, such measures could turn a growing drug resistance epidemic into a diminishing one (Fig.3B, blue vs green curves).

Nonetheless, in terms of overall TB burden, our results suggest that engaging the private sector alone will not be enough to meet the country’s aspirations for TB elimination. Rather, such measures lay the foundations for TB control by maximising the quality and coordination of basic TB services, across India’s vast and fragmented healthcare system ^16^. To explain limitations on PPSA transmission impact, our work highlights the complexity of the delay from symptoms to treatment initiation, showing how it arises from a combination of factors. For example, while the importance of diagnosis accuracy is well-recognised ^8,11,32^, pre-empting patient dropout, through offering a rapid diagnosis, can be as impactful for the diagnostic delay (Fig.3, Fig.6B – C). Second, our results suggest that the ‘patient delay’ in Fig.1A may play a larger role than previously recognized (Fig.4).

We note that this latter result is not directly measured, but inferred through reconciling ARTI and prevalence in the model. Previous studies have approached patient and diagnostic delays through retrospective patient interviews in various settings in India: a recent meta-analysis of these studies ^8^ found a median patient delay of around 18 days. To our knowledge there is no other independent, direct evidence for the ‘true’ patient delay. Nonetheless, there are some notable comparisons in a recent TB prevalence survey in Gujarat state. Of the bacteriologically positive TB cases, only 28% had sought care for their symptoms, including 11% that were on TB treatment ^33^. Although cross-sectional, these survey findings appear consistent with the picture of substantial transmission occurring independently of the ‘diagnostic delay’.

There are several possible reasons for these discrepancies between model and prevalence survey findings on the one hand, and patient interviews on the other. For example, in urban areas with poor air quality, prolonged cough is a common symptom: TB patients may tend to visit a provider when their symptoms become more advanced (e.g. fever), ultimately reporting only the duration of these more developed symptoms. Alternatively, the patient delay may truly be as short as 18 days, but only amongst those patients who seek care: there may remain a patient population who never contact the healthcare system, for example due to the opportunity costs of doing so. These factors may differ by region in India, as well as by gender and urban/rural setting. As illustrated by Fig.S4, mitigating these factors could maximize the impact of a PPSA.

Approaches towards mitigating these factors could involve active case-finding (ACF) ^34^. India’s recent National Strategic Plan underlines the potential importance of ACF in risk groups such as urban slums ^16^, while recent work in Viet Nam has also demonstrated the potential value of screening close contacts of diagnosed TB cases, together with longitudinal followup of these contacts ^35^. However, it is also possible for the patient delay to be impacted by measures to improve the demand for TB services; for example, social protection mechanisms ^36^ could have the secondary effects of encouraging TB symptomatics to come forward for care ^37^. Such effects are currently hypothetical, and present an important evidence gap for future studies to address.

As with any modelling approach, our model has several limitations to note. First, it takes a simplified view of the host population, essentially averaging over variations by gender and age. In future, better data on careseeking and the quality of care with respect to these factors would support a more refined approach incorporating these factors. Second, our work concentrates on two major cities in India, informed by the available, community-based studies on careseeking pathways. Further work, deploying such surveys more broadly, should explore to what extent these findings may be generalized to other cities India; one potentially important factor is the greater HIV burden in states like Andhra Pradesh ^3^. Moreover, this work does not address potential impact in rural settings. Indeed, recent work has highlighted the phenomenon of TB prevalence being higher in rural areas than urban ^38^, suggesting even longer delays before initiation of appropriate TB treatment: there is therefore a pressing need for a better understanding of healthcare utilisation in these settings. Third, we have made several simplifying assumptions on provider behaviour, namely that ‘engaged’ providers would show the same standard of care as in the public sector. As noted above, it is promising that the PPSA pilots have shown a dramatic increase in the number of TB cases being notified ^1,3^ ongoing data collection during the pilots will cast light on the extent to which the quality of TB care has been improved. Lastly, these results are quite sensitive to underlying assumptions about prevalence and ARTI, as well as to transmission over the course of illness. If more transmission is occurring at later stages of illness, then private provider engagement could more effectively interrupt transmission and avert twice as many cases as our baseline uninformed assumption of uniform infectivity. Objective data on the ‘transmission curve’ would be useful to clarify the appropriate baseline for these and most TB models.

In summary, private sector engagement is a key foundation for managing TB in India. In addition to its direct benefit to TB patients, an engaged private sector will also enable the maximum deployment of future interventions against TB in India. While building such favourable conditions for TB control, there is an urgent need to identify where TB transmission is occurring: only by addressing this transmission will it truly be possible to accelerate declines in India’s vast TB epidemic.

## Additional information

The authors have no competing interests that might be perceived to influence the results and/or discussion reported in this paper.

## Author contributions

NA, PD conceived the study. NA, SD, SS performed the analysis; SK, RR, BV, NK, DG, KR, SAN and PD contributed to the interpretation and validation of model results; NA wrote a first draft of the manuscript, and all authors contributed to the final draft.

## Supporting information

### 1. Model specification

The model is governed by the following equations (see table S1 for definitions of state variables, and table S2 for parameter definitions and sources). First, for the states prior to a TB patient’s first visit to a provider, we have:

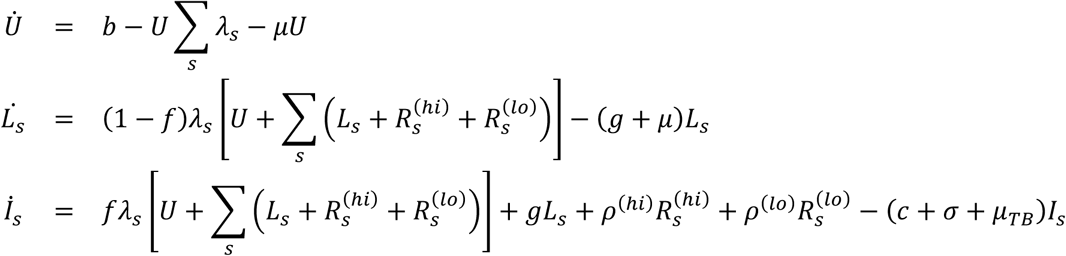

where the subscript s denotes the infecting strain (with values 0,1 denoting drug-susceptible and drug resistant TB, respectively). The parameter *c* is the rate of careseeking, its inverse representing the average patient delay before first presentation for care.

Upon presenting for care, we assume that a proportion *p_r_* of patient visit a provider of type *r* (denoting the public sector; FQ providers; LTFQ providers; and chemists – see table S1). We have, for those awaiting diagnosis with provider type *r* and infected with strain s:

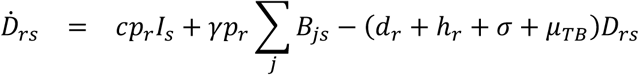

As described in the main text, a patient-provider interaction may last days to weeks. This stage ends either when the provider finally offers a diagnosis (whether correctly for TB or otherwise), or when the patient leaves the provider, to seek care elsewhere. Here, we model these two endpoints through competing hazards of offering a diagnosis (*d_r_*), versus the patient leaving the provider (*h_r_*). As described below, these rates are estimated from the patient pathway surveys conducted in Mumbai and Patna ^1^.

We assume that a proportion *u_r_* of TB patients visiting provider type *r* successfully initiate TB treatment (the remainder constituting missed diagnosis as well as initial loss to followup, covered below). For those initiating first-line treatment, it is convenient to specify equations separately by drug-susceptible (*s* = 0) and drug-resistant (*s* = 1) status. Thus we have, for drug-susceptible TB:

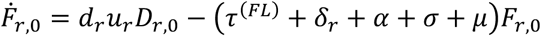

where *δ_r_* is the per-capita rate of default from first-line treatment with provider type *r* and *α* represents the per-capita hazard of acquisition of multi-drug-resistance while on first-line TB treatment, only applicable to drug-sensitive TB. We assume that those defaulting from treatment are bacteriologically negative, but have an elevated risk of relapse, in comparison with those who have successfully completed treatment. The relevant compartments are discussed below.

For drug-resistant TB on first-line treatment, we have:

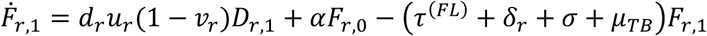

where *v_r_* is the proportion of TB patients presenting to a provider of type *r* who undergo drug sensitivity testing at the point of TB diagnosis.

For second-line treatment (only for DR-TB), we have:

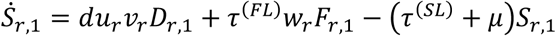

where *w_r_* represents the proportion of DR-TB patients with provider type *r* who are switched to second-line treatment after failing first-line treatment.

Next, the compartment *B* captures those patients who have dropped out of the care cascade and remain infectious, whether by failed diagnosis, loss to follow up, or failed treatment. We have, for *B*:

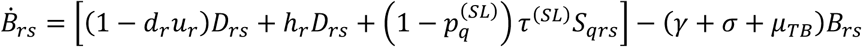

Those who have recovered from disease include patients who have completed treatment; those who have defaulted from treatment; and those who have recovered spontaneously. We assume the latter two to have an elevated risk of relapse compared to the former, in the two years following recovery. Following this period, remaining recovered individuals stabilize in their relapse risk. Thus we have, for the ‘high’ and ‘low’ relapse risk compartments, respectively:

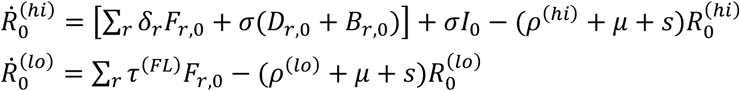

Finally, for the forces-of-infection λ_0_, λ_1_ for DS- and DR-TB respectively, we have:

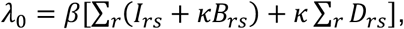

and likewise for λ_1_, but with **β*_MDR_* in place of *β*.

### 2. Patient pathways

We used data from community-based patient pathway surveys, recently conducted in Mumbai (76 TB patients and 196 patient-provider interactions) and Patna (64 TB patients and 121 patient-provider interactions), and described in detail elsewhere ^1^. In brief, individuals in the community, who had been on TB treatment within the preceding 6 months, were administered an in-depth interview, to identify the sequence and types of providers that each patient visited before their TB diagnosis.

A patient’s contact with a given provider may last several days, sometimes weeks: this process ends either when the provider eventually makes a diagnosis, or when the patient drops out to visit an alternative provider. Here, we model this combination of behaviours using independent, competing exponential hazards with rates *r_Diagnosis_* and *r_Dropout_*, specific to the type of provider involved (public, FQ, LTFQ or chemist). Figure 1B shows the overall framework: for Mumbai and Patna separately. The rates are estimated from the data and their reciprocals give us the average time of diagnosis and the average time of dropout, respectively, for each type of provider. The probability of getting a diagnosis at a provider (whether a correct diagnosis or not) is equal to 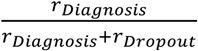, and we estimate the accuracy of diagnosis of each type of provider from the data. We also model the role of different provider types in the careseeking pathway, in particular: the proportions of patients visiting each type of provider on the first careseeking attempt, and the corresponding proportions on subsequent visits, conditional on the type of provider last seen. 30 of the 196 patient-provider interactions in Mumbai, and 11 of the 121 patient-provider interactions in Patna, are such that the providers consulted are private, however, their qualifications, and hence their types (LTFQ/FQ), are missing. We let each missing provider type be represented by an unknown binary variable. Since the model parameters are specific to the provider type, the expression for the likelihood of the pathways data as a function of the model parameters also involves the missing binary variables. We use the iterative algorithm *Expectation Maximization* (EM) to obtain the maximum likelihood estimates of parameters. Each iteration involves two steps: *E-Step:* Finding the expectation of the log likelihood function, over the distribution of the missing binary variables conditioned on the observed data, under an initial estimate of the parameters, and *M- Step:* Maximizing the expectation of the log likelihood function to obtain a revised estimate of parameters. The revised estimate is then used as an initial estimate for the E-Step, and the process continues until the values of the maximum expectation of the log likelihood converge within a specified tolerance. The associated variance-covariance matrix of the estimates is approximated as the inverse of the observed Fisher Information Matrix, which is equal to the difference of the negative of the expectation of the Hessian matrix of the complete data log likelihood function, conditioned on the incomplete data and the expectation of the square of gradient of complete data log-likelihood function, conditioned on the incomplete data; evaluated at the final iteration of the EM algorithm.

For parameters related to the treatment cascade (the proportion of TB diagnoses initiating and completing treatment), we draw from a recent systematic review for the public sector ^2^. In the absence of systematic evidence for private providers, we incorporate plausible uncertainty distributions for these parameters (Table S2).

### 3. Model calibration and propagating uncertainty

We denote by *θ* the vector of input parameters, including *β*, *β*_*MDR*_, *c*, and other model inputs subject to uncertainty. For a given country, and a given parameter set *θ*, we initially simulated the model to equilibrium in the absence of the public sector (e.g. as in ref. ^3^) and MDR-TB, to capture the early history of the TB epidemic. Subsequently allowing population growth, we initiated the emergence and spread of DR-TB from 1980. We also captured the expansion of the public sector as a linear increase in *p*_0_ during the years of RNTCP scale-up, i.e. from 1997 to 2006 ^4^. By combining these processes, we determined model projections for calibration targets (prevalence, ARTI and percent of incident TB cases being drug-resistant), assumed to apply in 2017.

To compare these model projections with data *D*, we defined the *posterior density π(θ)* as:

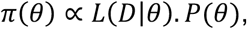

where *L* is the likelihood of the data *D* given θ and *P* is the joint prior distribution for θ. For P, we took independent uniform distributions over the ranges shown in table S2 (taking +/− 20% of the point values where no ranges are shown). The likelihood *L* is constructed as follows.

We fitted a log-normal distribution to prevalence: in particular, we determined the mean and variance of this distribution in order for the 2.5^th^, 50^th^ and 97.5^th^ percentiles to match respectively the lower, mid and upper ranges of prevalence estimates. We write *F*^(*Prev*)^(·) for the probability density thus obtained. Likewise, we write *F*^(*ARTI*)(·), *F*(*pMDR*)^(·) for the inferred probability densities corresponding respectively to prevalence and the proportion of incident TB that is MDR in year *t*. Then we have, for the overall likelihood:

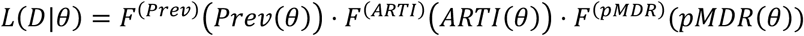

where, for example, *ARTI(θ)* is the simulated value of incidence in 2017 given parameters *θ*, and likewise for the other functions of *θ* in the expression above. In practice we compute the logarithm of π(*θ*), thus taking the sum of the logarithms of each of the terms shown above.

With *π(*θ*)* thus defined, we sampled the posterior density using a Markov Chain Monte Carlo approach. In brief, this approach implements a random walk through the space of parameter values I to obtain an unbiased sample of the posterior density. We implemented the ‘adaptive’ MCMC algorithm first introduced by Haario et al ^5^, which incorporates a dynamic covariance matrix to adjust endogenously the scale of ‘jumps’ in proposals for each of the parameter values. For the set of parameter values thus obtained, we took every tenth element to reduce autocorrelation, thus yielding an ‘ensemble’ of parameters *θ*_1_, *θ*_2_,… This ensemble captures simultaneously the uncertainty in the parameter inputs, as well as in the calibration data. Then, to estimate uncertainty in given simulated outputs (e.g. in the reduction of incidence with a given coverage of intervention), we simulated this output *Γ_i_* for every **θ**_*i*_. We finally estimated uncertainty in *Γ_i_* by determining its 2.5^th^, 50^th^ and 97.5^th^ percentiles.

**Figure S1.**
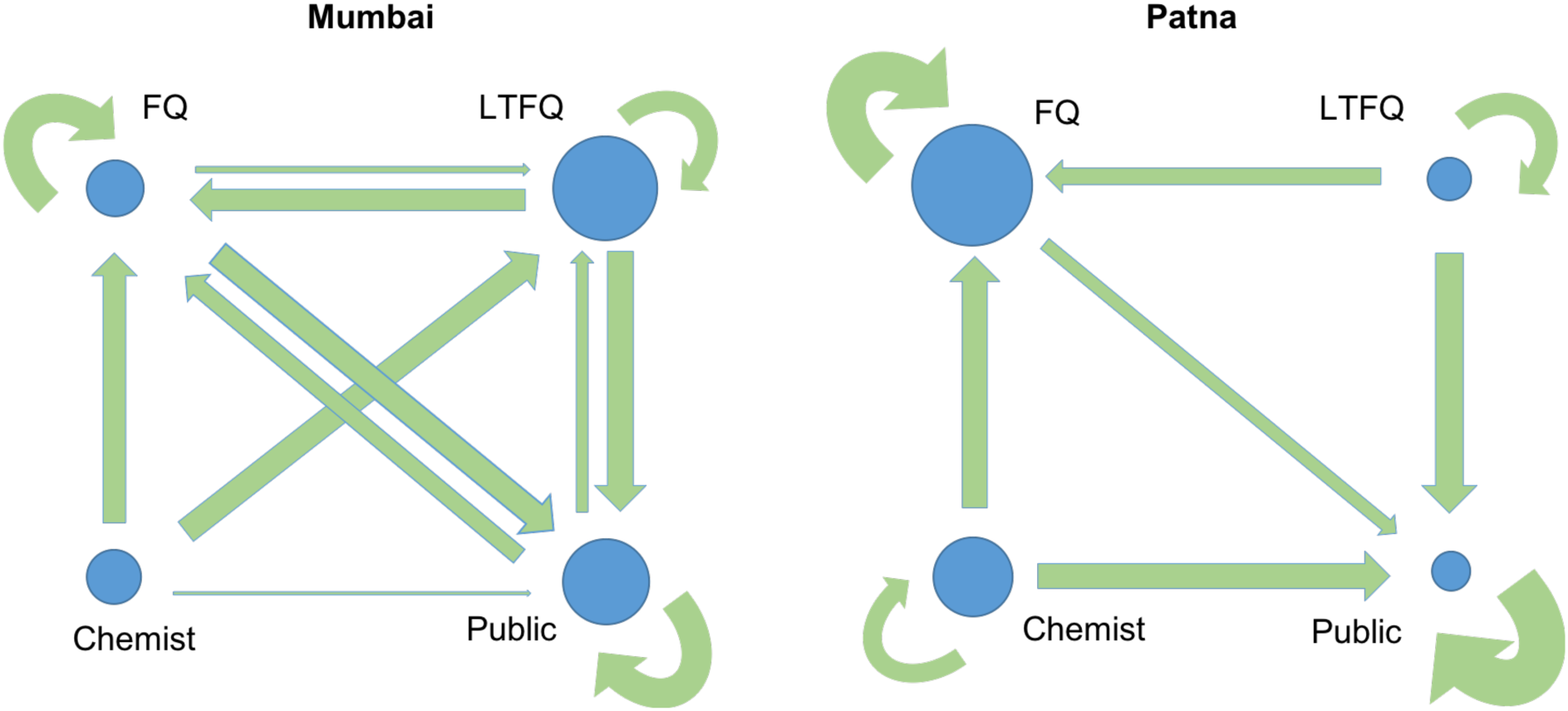
Summary of the contrasting patient careseeking pathways in Mumbai and Patna. Circle areas are proportional to the importance of providers as first point of patient contact (for example, patients in Patna tend to seek care first amongst fully qualified providers). Arrows denote how patients switch providers on subsequent visits, with arrow widths proportional to frequency.

**Figure S2.**
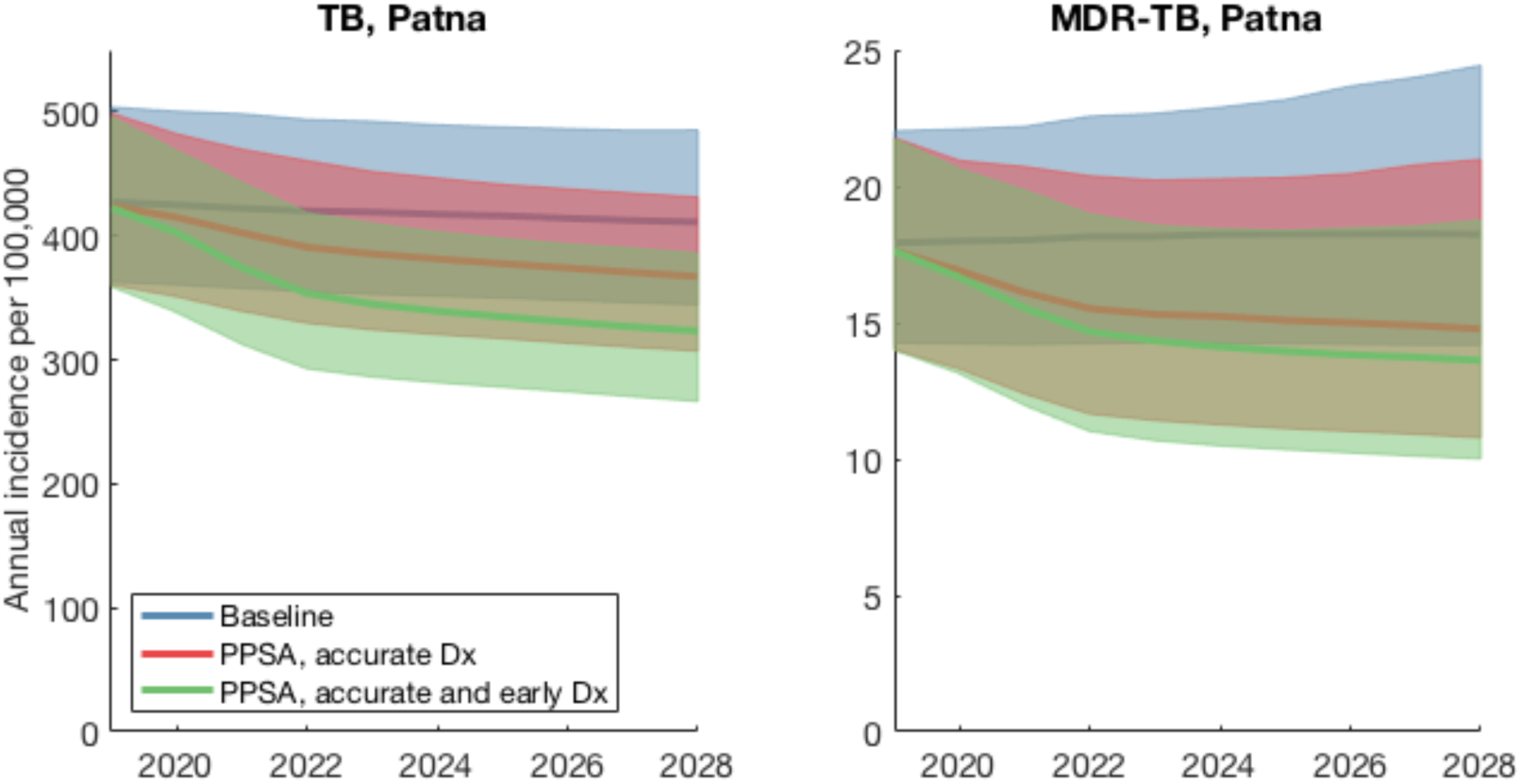
Simulated impact of a PPSA in Patna. As for Figure 3 in the main text, but for Patna. See Figure 3 caption for further details.

**Figure S3.**
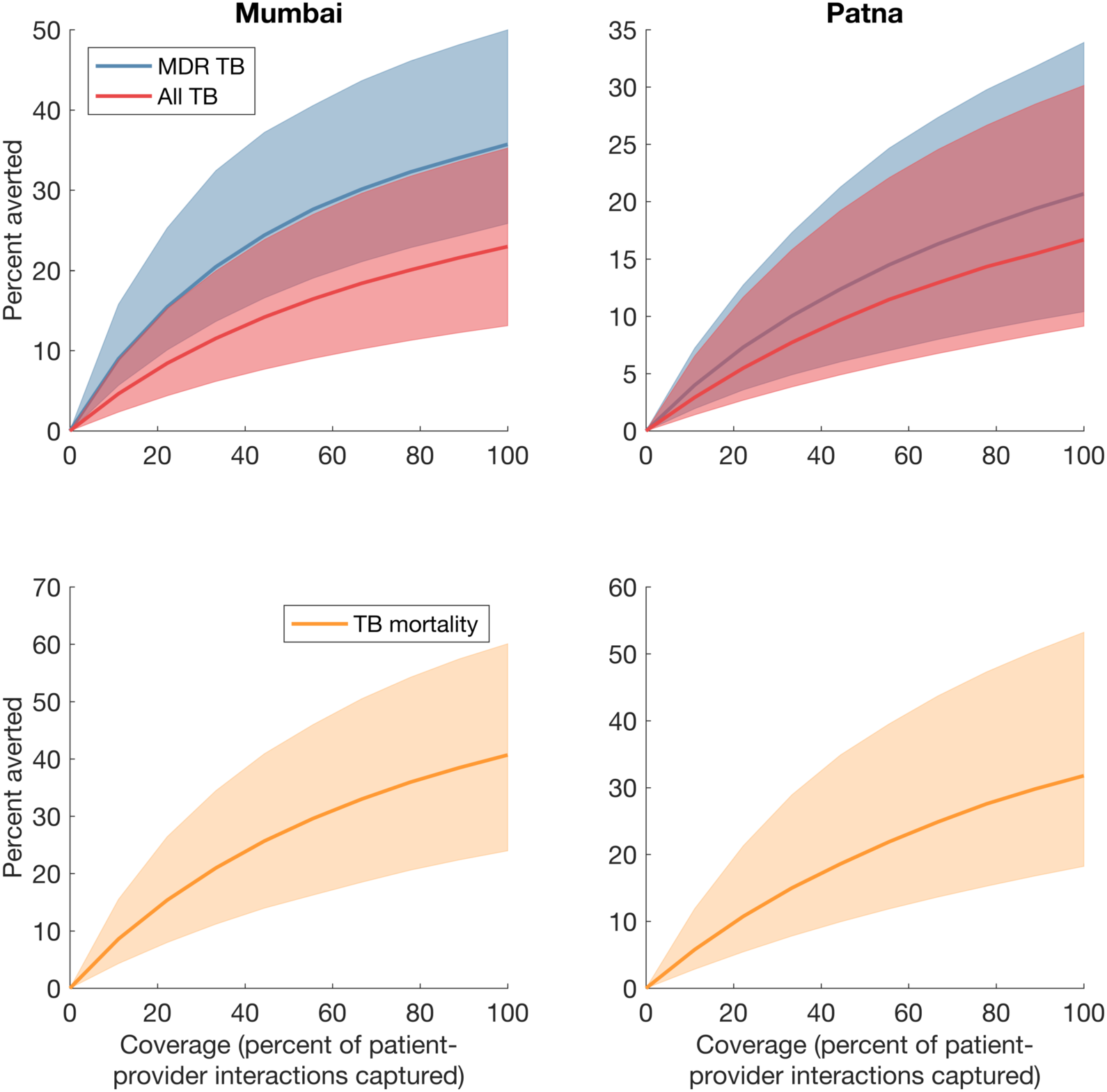
**Potential impact of a PPSA at different levels of coverage**, in Mumbai (left-hand column) and in Patna (right-hand column). Lines show central estimates, and shaded regions show 95% credible intervals.

**Figure S4:**
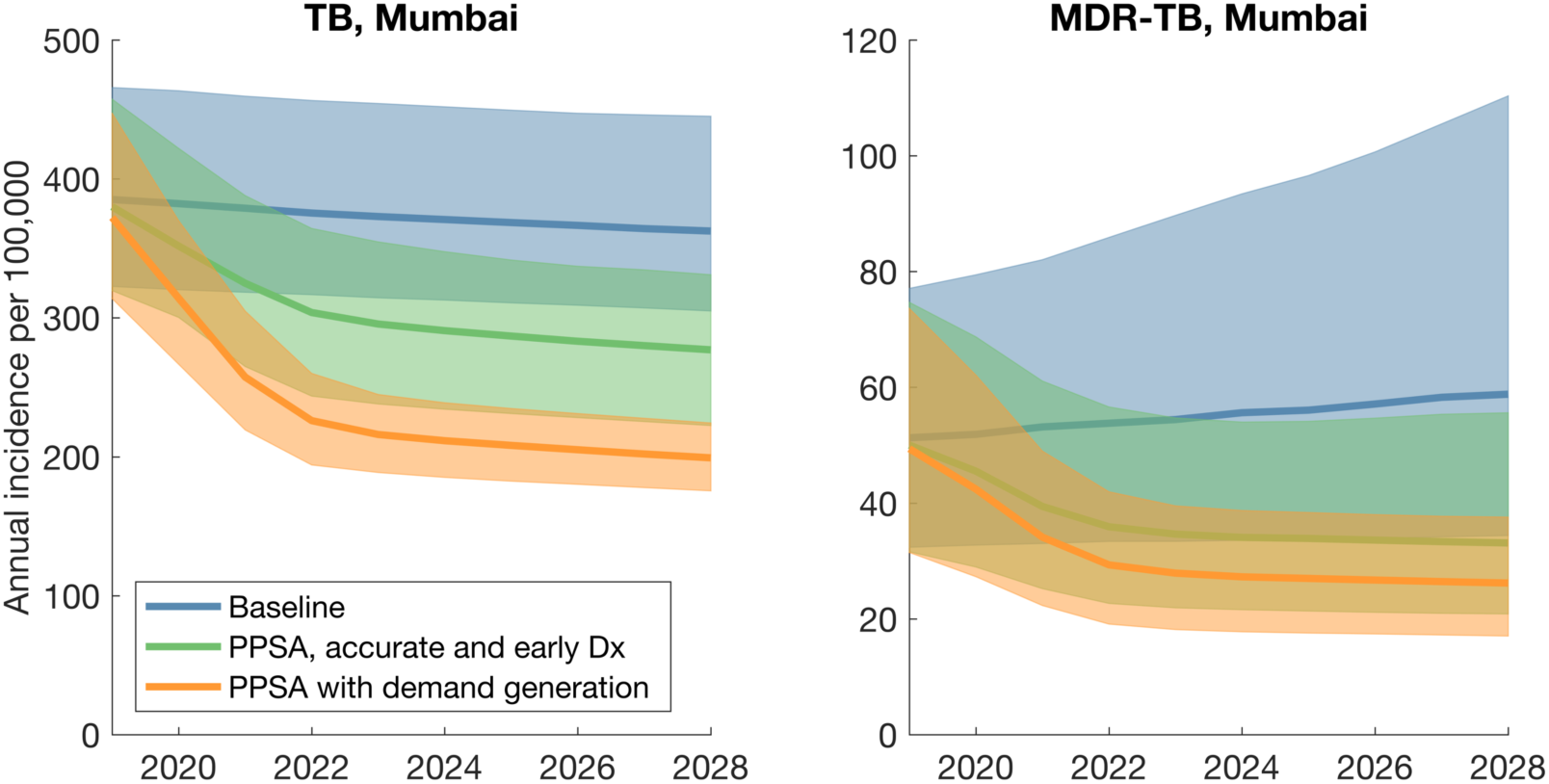
**Potential PPIA supplemented by demand generation** (yellow curve). This provides the same scenarios shown in Fig.3 in the main text (a PPSA in Mumbai at 75% scale), but with ‘demand generation’ added for comparison. Uncertainty regions not shown, for clarity. Here, demand generation is assumed to bring about a 40% reduction in the patient delay. Such measures could involve lowering the barriers for access to care, or intensified case-finding. The impact shown here corresponds to a 37% (95% CrI 30.9 – 43.8%) reduction in cumulative incidence.

**Table S1.**
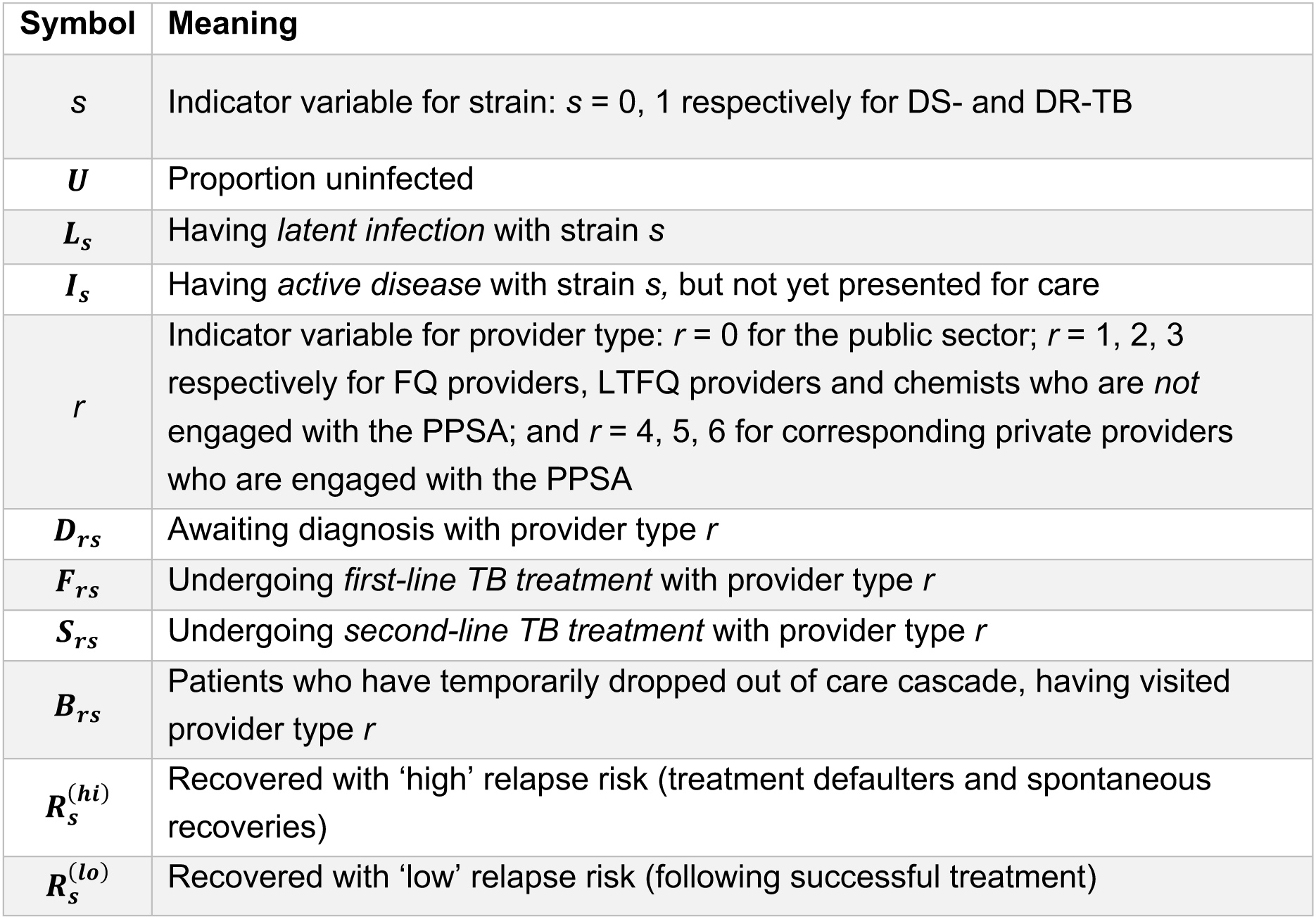
List of state variables used in the model.

| Symbol | Meaning |
| --- | --- |
| $s$ | Indicator variable for strain: $s = 0, 1$ respectively for DS- and DR-TB |
| $U$ | Proportion uninfected |
| $L_s$ | Having *latent infection* with strain $s$ |
| $I_s$ | Having *active disease* with strain $s$, but not yet presented for care |
| $r$ | Indicator variable for provider type: $r = 0$ for the public sector; $r = 1, 2, 3$ respectively for FQ providers, LTFQ providers and chemists who are *not* engaged with the PPSA; and $r = 4, 5, 6$ for corresponding private providers who are engaged with the PPSA |
| $D_{rs}$ | Awaiting diagnosis with provider type $r$ |
| $F_{rs}$ | Undergoing *first-line TB treatment* with provider type $r$ |
| $S_{rs}$ | Undergoing *second-line TB treatment* with provider type $r$ |
| $B_{rs}$ | Patients who have temporarily dropped out of care cascade, having visited provider type $r$ |
| $R_s^{(hi)}$ | Recovered with 'high' relapse risk (treatment defaulters and spontaneous recoveries) |
| $R_s^{(lo)}$ | Recovered with 'low' relapse risk (following successful treatment) |

**Table S2.** Input parameters and data used for the model.

| Parameter | SYMBOL |  | VALUE | SOURCE/COMMENTS |
| --- | --- | --- | --- | --- |
| TB epidemiology |  |  |  |  |
| Annual risk of TB infection (ARTI) in urban slums as of 2015 | | $\lambda$ | 2 – 3% | Gopi (2008)<sup>6</sup> |
| TB prevalence in urban slums as of 2015 | | $P$ | 388 per 100,000 (233 – 543) | Baskaran (2015)<sup>7</sup> |
| Percent of incident TB cases that are MDR | Mumbai | $p_{\text{MDR}}$ | 12% (8 – 16) | Assumption |
| | Patna | $p_{\text{MDR}}$ | 4% (3 – 5) | Assumption |
| TB natural history |  |  |  |  |
| Per-capita rate of reactivation of latent TB | | $r$ | 0.001 yr-1 | Horsburgh (2010)<sup>8</sup> |
| Proportion of infections undergoing rapid progression | | $p_{fast}$ | 0.1 | Vynnycky (1997)<sup>9</sup> |
| Per-capita rate of relapse | Low relapse risk | $\rho_{lo}$ | 0.002 yr-1 (0.001 - 0.004) | Menzies (2009)<sup>10</sup> |
| | High relapse risk | $\rho_{hi}$ | 0.02 yr-1 (0.01 - 0.04) | |
| Per-capita mortality hazard | Non-TB | $\mu$ | 0.0152 yr-1 | World Bank estimates |
| | TB cases | $\mu_{TB}$ | 0.089 yr-1 (0.33 - 1.21) | Tiemersma (2011)<sup>11</sup> (averaged across smear-positive and smear-negative TB), corresponding to 50% mortality in an average of 3 years |
| Per-capita rate of spontaneous cure | | $\sigma$ | 0.089 yr-1 (0.33 - 1.21) | |
| Anti-TB treatment (a, b) |  |  |  |  |
| Rate of completion of first-line TB treatment | | $\tau_1$ | 2 yr-1 | Corresponds to a duration of 6 months |
| Rate of completion of second-line TB treatment | | $\tau_2$ | 0.5 yr-1 | Corresponds to a duration of 2 years |
| Proportion of diagnosed TB cases initiating first-line TB treatment | Public | $p_{pu}^{FLinit}$ | 90% | RNTCP (2015) <sup>12</sup> |
| | Private | $p_{pr}^{FLinit}$ | 0.6 (0.4 - 0.8) | Assumption |
| Proportion of diagnosed TB cases initiating second-line TB treatment | Public | $p_{pu}^{SLinit}$ | 90% | RNTCP (2015) <sup>12</sup> |
| | Private | $p_{pr}^{SLinit}$ | -- | |
| Proportion completing first-line TB treatment | Public | $p_{pu}^{FLcomp}$ | 85% | RNTCP (2015) <sup>12</sup> |
| | Private | $p_{pr}^{FLcomp}$ | 0.6 (0.4 - 0.8)% | Uplekar (1998) <sup>13</sup> |
| Proportion completing second-line TB treatment | Public | $p_{pu}^{SLcomp}$ | 0.5 | RNTCP (2015) <sup>12</sup> |
| | Private | $p_{pr}^{SLcomp}$ | -- | |
| Per-capita rate of acquiring multi-drug resistance while on first-line treatment | Public | $m_{pu}$ | 0.01 yr-1 | Menzies (2009) <sup>10</sup> |
| | Private | $m_{pr}$ | 0.05 yr-1 | |
| Proportion of MDR-TB cases receiving drug susceptibility testing at point of TB diagnosis | Public sector | $dst_{pu}$ | 0.15 (0.05, 0.25) | Using data from national GeneXpert demonstration<sup>14</sup> |
| | Private (any type) | $dst_{pr}$ | 0 | Assumption (c) |

Notes:

**Table S3.** City-specific pathway parameters, inferred from the pathway data.

|  |  | Public | FQ | LTFQ | Chemists |
| --- | --- | --- | --- | --- | --- |
| **Mumbai** |  |  |  |  |  |
| *Proportion visited on initial consultation* |  | 0.30 (0.15, 0.47) | 0.13 (0.06, 0.22) | 0.45 (0.33, 0.57) | 0.12 (0.06, 0.19) |
| *Proportion visiting after leaving provider of type:* | Public | 0.47 (0.24, 0.75) | 0.32 (0.14, 0.53) | 0.21 (0.06, 0.39) | 0 (0, 0) |
|  | FQ | 0.46 (0.16, 0.81) | 0.43 (0.19, 0.72) | 0.11 (0, 0.3) | 0 (0, 0) |
|  | LTFQ | 0.46 (0.23, 0.70) | 0.39 (0.20, 0.58) | 0.15 (0.03, 0.29) | 0 (0, 0) |
|  | Chemists | 0.11 (0, 0.57) | 0.44 (0.17, 0.77) | 0.44 (0.17, 0.77) | 0 (0, 0) |
| *Probability of TB diagnosis per provider visit* |  | 0.94 (0.91, 0.98) | 0.93 (0.88, 0.98) | 0.75 (0.56, 0.94) | 0 (0, 0) |
| *Hazard Rate, provider offering diagnosis* |  | 0.087 (0.069, 0.106) | 0.074 (0.052, 0.096) | 0.035 (0.019, 0.051) | 0 (0, 0) |
| *Hazard rate, patient shopping* |  | 0.04 (0.03, 0.05) | 0.04 (0.03, 0.06) | 0.05 (0.03, 0.07) | 0.06 (0.05, 0.08) |
| **Patna** |  |  |  |  |  |
| *Proportion visited on initial consultation* |  | 0.063 (0, 0.29) | 0.61 (0.46, 0.77) | 0.08 (0, 0.20) | 0.25 (0.14, 0.36) |
| *Proportion visiting after leaving provider of type:* | Public | 1 (1, 1) | 0 (0, 0) | 0 (0, 0) | 0 (0, 0) |
|  | FQ | 0.24 (0.08, 0.42) | 0.76 (0.58, 0.95) | 0 (0, 0) | 0 (0, 0) |
|  | LTFQ | 0.5 (0, 1) | 0.3 (0, 0.9) | 0.20 (0, 0.69) | 0 (0, 0) |
|  | Chemists | 0.44 (0.22, 0.72) | 0.44 (0.22, 0.67) | 0 (0, 0) | 0.11 (0, 0.26) |
| *Probability of TB diagnosis per provider visit* |  | 1 (1, 1) | 0.95 (0.89, 1) | 0 (0, 0) | 0 (0, 0) |
| *Hazard Rate, provider offering diagnosis* |  | 0.18 (0.14, 0.23) | 0.10 (0.07, 0.14) | 0.0 (0, 0.17) | 0 (0, 0) |
| *Hazard rate, patient shopping* |  | 0.008 (0.006, 0.011) | 0.06 (0.04, 0.08) | 0.14 (0.04, 0.27) | 0.12 (0.09, 0.16) |

**Table S4.** Model outputs for key transmission parameters.

|  | Infectivity, mean infections per year per case |  | Mean duration, patient delay (months) |
| --- | --- | --- | --- |
| | $\beta_{DS}$ | $\beta_{MDR}$ | $1/d$ |
| **MUMBAI** | 16.1 (9.7 – 28.8) | 12.8 (7.9 – 20.9) | 4.39 (1.64 – 9.57) |
| **PATNA** | 15.1 (10.1 – 27.8) | 8.9 (5.3 – 17.5) | 5.24 (2.19 – 9.40) |

